# SMOC1 Regulates Endothelial-to-Mesenchymal Transition During Cardiac Repair After Myocardial Infarction

**DOI:** 10.64898/2025.12.10.693412

**Authors:** Fatima Hacer Kurtoglu Babayev, Bardia Amirmiran, Fredy Delgado Lagos, Beate Fisslthaler, Ralf P. Brandes, Roman Vuerich, Mauro Siragusa, Ingrid Fleming

**Author notes:** Address correspondence to: Ingrid Fleming PhD. Goethe University Frankfurt, Institute for Vascular Signalling, Centre for Molecular Medicine, Theodor Stern Kai 7, 60596 Frankfurt am Main, Germany.

## Abstract

Endothelial-to-mesenchymal transition (EndMT) is a crucial, dual-phase process in cardiac repair after myocardial infarction (MI), driving both initial scar stabilization and subsequent pathological fibrosis. Therapeutic targeting requires precise temporal control rather than complete inhibition. This study identifies the matricellular protein SPARC-related modular calcium-binding protein 1 (SMOC1) as a novel regulator of EndMT. Analysis of single-cell RNA sequencing data from post-MI mouse hearts revealed that SMOC1 is highly enriched in a subpopulation of endothelial cells undergoing late EndMT. In vitro, SMOC1 expression was upregulated during cytokine-induced EndMT in human endothelial cells. Its siRNA-mediated knockdown exacerbated the EndMT phenotype, increasing mesenchymal marker expression and cell morphology changes, effects rescued by recombinant SMOC1 (rSMOC1). Mechanistically, SMOC1 deficiency enhanced TGF-β2-induced SMAD2 phosphorylation, while rSMOC1 attenuated this pathway and promoted a shift from the short to the long, signaling-competent isoform of endoglin. In vivo, endothelial-specific SMOC1 deficiency (SMOC1^ΔEC^) in mice promoted age-associated EndMT and profoundly worsened post-MI outcomes. After MI, SMOC1^ΔEC^ mice exhibited exacerbated cardiac dysfunction, ventricular dilation, pathological fibrosis, increased inflammatory cell infiltration, reduced survival, and a higher incidence of cardiac rupture compared to controls. Collectively, these findings establish SMOC1 as a critical endogenous modulator of EndMT that restrains its pathological progression. SMOC1 coordinates endothelial cell phenotype, in part by fine-tuning TGF-β/endoglin signaling, and its loss accelerates maladaptive remodeling post-MI. Thus, SMOC1 represents a potential therapeutic target for spatially and temporally controlling EndMT to improve cardiac repair

## Introduction

Endothelial-to-mesenchymal transition (EndMT) is a critical cellular reprogramming process in which endothelial cells change their phenotype and adopt more mesenchymal cell-like characteristics.^1^ It plays a key role in cardiac repair after myocardial infarction (MI), characterized by dual protective and pathological effects.^2,3^ In the acute phase, driven by hypoxia and transforming growth factor (TGF)-β signaling, EndMT helps form the fibroblasts and myofibroblasts that contribute to the formation of a stabilizing scar, preventing ventricular rupture.^4–6^ However, the persistence of EndMT becomes a major driver of adverse remodeling and heart failure. Sustained EndMT leads to excessive deposition of stiff extracellular matrix (ECM), contributing to pathological fibrosis that impairs cardiac compliance and function.^7^ This fibrotic process extends beyond the initial infarct zone into remote myocardium, promoting ventricular stiffening and arrhythmogenesis. A particularly detrimental consequence is the disruption of vascular integrity and repair. As endothelial cells transdifferentiate, they deplete the vascular pool, directly impairing angiogenesis and capillary density.^8,9^ This creates a vicious cycle where ischemia perpetuates further EndMT and fibrosis. Beyond post-ischemic remodeling, emerging evidence suggests that chronic, low-grade EndMT contributes to progressive deposition of ECM. This results in the vascular stiffening and endothelial dysfunction that characterize the aged myocardium and vasculature. Consistently, mesenchymal gene programs are upregulated in multiple tissues during aging, particularly in endothelial cells.^10^

EndMT represents a key therapeutic target but the therapeutic goal would not be indiscriminate suppression of the process, as this would compromise early wound stability. Rather, interventions should be aimed at the temporal and spatial control of the process. Therefore, this study set out to identify novel markers of EndMT and assess their role in the response to acute damage as well as post-infarct remodeling.

## Results

### SMOC1 is enriched in the EndMT subpopulation of cardiac endothelial cells after MI

The analysis of single cell transcriptomes from cardiac tissue up to 28 days after MI^6^ revealed a significant impact of infarction on several cell types including macrophages, fibroblasts B cells and endothelial cells (**Fig. 1A**). A distinct subpopulation of endothelial cells (cluster 2) emerged after MI that expressed markers of late EndMT including the ECM and mesenchymal transition genes *Col1a1, Col3a1, Fn1, Acta2, Tagln, Vim, Postn, Timp1, Serpine1, Mmp2, Itgb1, Cdh2, Thbs1, Fbln2*, and *Lox* (**Fig. 1B-C**). This cluster also expressed *Smoc1*, which encodes a secreted matricellular protein.^11,12^ Although *Smoc1* was detectable in fibroblasts and in endothelial cells undergoing early EndMT, its expression was markedly enriched in the late EndMT cluster, with elevated levels persisting up to day 7 after MI (**Fig 1D**). Indeed, SMOC1 levels in cardiac endothelial cells showed an inverse relationship with that of CD144 (Cdh5) and increased and then decreased in parallel with fibronectin (FN1) (**Fig. 1D**). Thus SMOC1 was temporally associated with a transient EndMT phenotype, or endothelial to mesenchymal activation^6^ after MI.

**Fig. 1.**
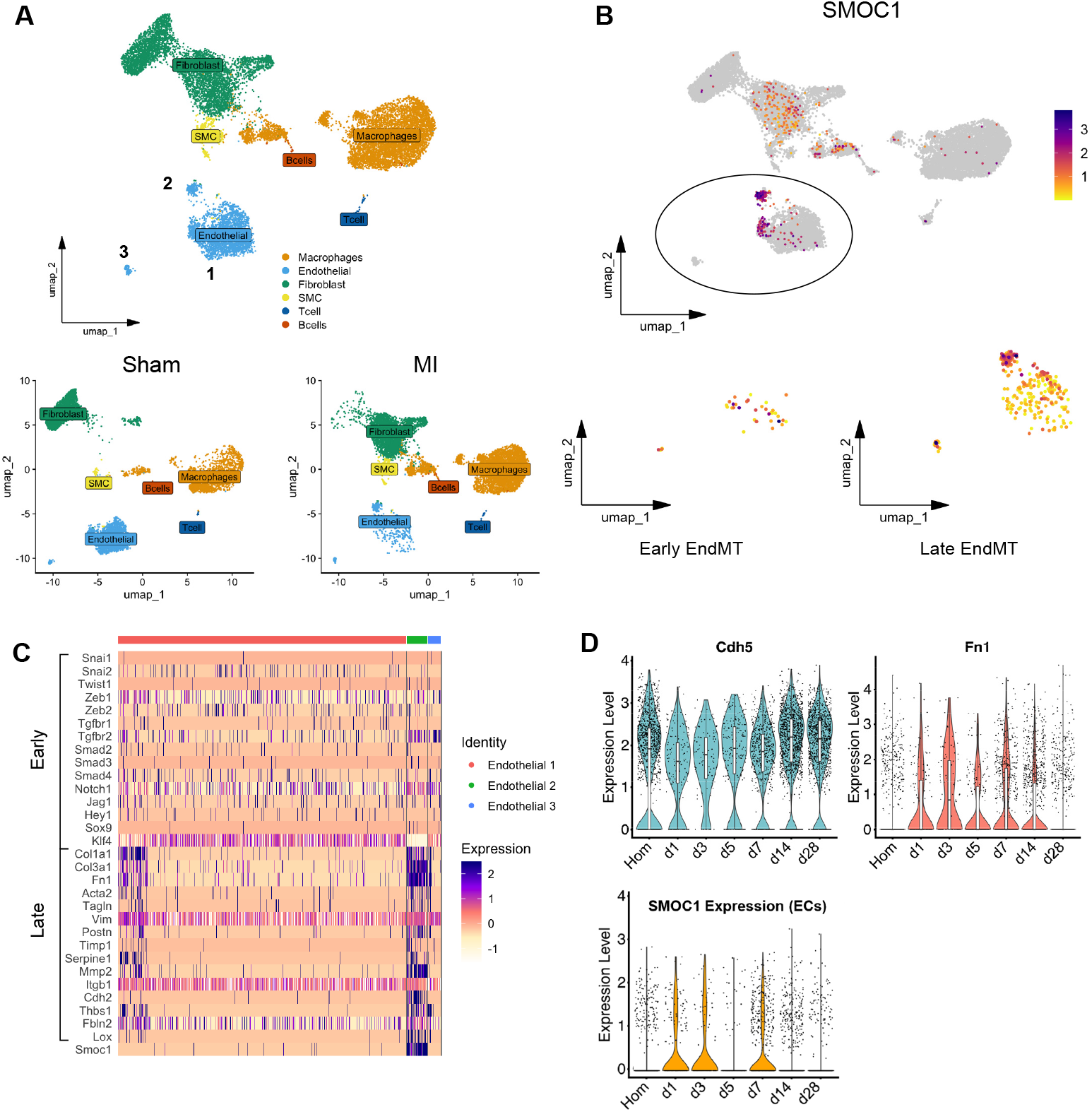
Relationship between the expression of EndMT markers and *SMOC1* after acute myocardial infarction (**A**) scRNA-seq UMAP projection showing cardiac cell types isolated from 13 mice after MI. Colours represent major cell lineage clusters, including endothelial cells, macrophages, fibroblasts, smooth muscle cells (SMC), T cells and B cells. The lower panels show cell type distribution in sham-treated and MI hearts. (**B**) SMOC1 expression in the cell clusters outlined in A. The lower panels show refined clustering analysis of the endothelial cell populations. (**C**) Heatmap showing gene expression changes in the three endothelial cell clusters identified in A and classification as early and late EndMT. (**D**) Violin plot showing the normalized expression level of *Smoc1* transcript specifically within the endothelial cell population at homeostasis (Hom) and up to 28 days after MI. Data points represent individual cells.

Next, EndMT was induced in endothelial cells using a combination of IL-1β and TGF-β2 and changes in gene expression were assessed by RNA sequencing. Consistent with previously published reports,^6^ EndMT induced an early loss of endothelial identity, marked by reduced expression of CD31 and increased expression of extracellular-matrix–associated genes such as FN1 (**Fig. 2A&B**). Gene ontology enrichment demonstrated activation of matrix remodeling, cytoskeletal dynamics and leukocyte migration programs expected with EndMT induction (**Fig. 2C**). As was the case in the *in vivo* dataset, SMOC1 was one of the genes significantly upregulated by EndMT as was endoglin (ENG, CD105), which we previous reported to bind SMOC1.^13^ The increase in SMOC was confirmed at the protein level in endothelial cells (**Fig. 2D**).

**Fig. 2.**
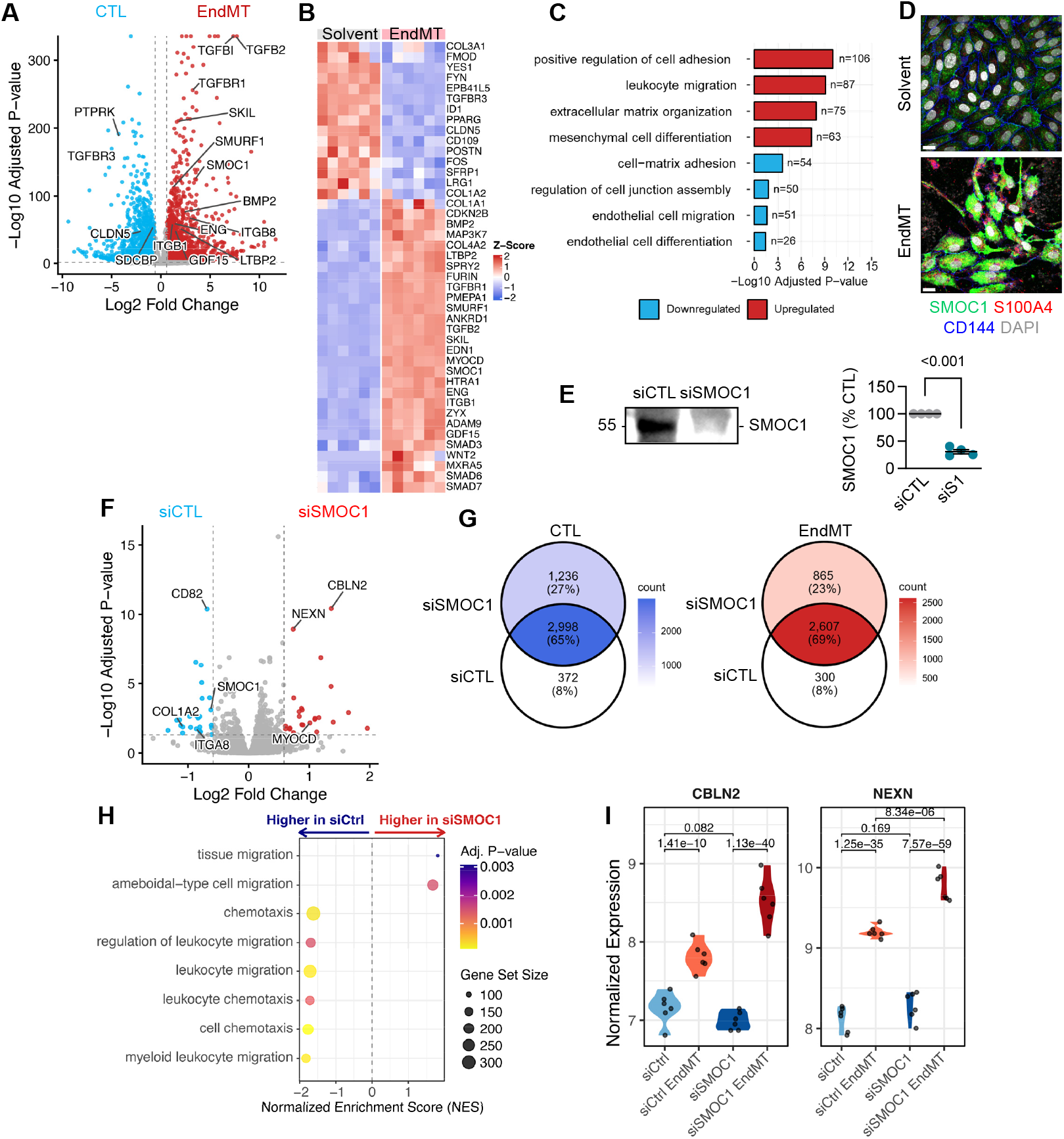
Effect of EndMT on SMOC1 expression and SMOC1 on EndMT. (**A**) Volcano plot showing differentially expressed genes (DEGs) in endothelial cells under control conditions (CTL) versus EndMT. Significantly altered DEGs (log2 Fold Change ≥ 0.585 and –log10 Adjusted P-value ≥ 1.3) are shown in red and blue; n= 6 independent cell batches. (**B**) Heatmap of the most altered DEGs in A. Color intensity represents Z-score normalized expression. (**C**) Pathway enrichment analysis (Over-representation analysis) showing the pathways most affected by EndMT induction. (**D**) Impact of EndMT on SMOC1, CD144 and S100A4 protein levels in endothelial cells; n=5 independent cell batches. (**E**) Effectiveness of the siRNA-mediated downregulation of SMOC1 versus a control siRNA (siCTL); n=4 independent cell batches. Student’s t-test. (**F**) Volcano plot showing the impact of SMOC1 downregulation (siSMOC1) versus a control siRNA on gene expression 72 hours after EndMT induction; n= 6 independent cell batches. (**G**) Venn diagrams showing the impact of siRNA-mediated SMOC1 deletion on endothelial cell gene expression under basal conditions (CTL) and 72 hours after EndMT induction. (**H**) GSEA of the genes altered by SMOC1 deletion versus a control siRNA (siCTL) 72 hours after EndMT induction. (**I**) CBLN2 and NEXN genes are consistently upregulated by siSMOC1 treatment in both baseline and EndMT conditions.

To determine whether SMOC1 was a marker of EndMT or could affect its development, we studied the impact of its deletion on gene expression in endothelial cells under control conditions and following the application of IL-1β and TGF-β2. The siRNA-mediated downregulation of SMOC1 efficiently abrogated protein expression (**Fig. 2E**) and induced marked alterations in the late EndMT transcriptome (**Fig. 2F**). The lack of SMOC1 affected the expression of 1165 genes induced by the cytokine cocktail, increasing the expression of 865 and decreasing the expression of 300 versus cells treated with a control siRNA (**Fig. 2F**), affecting largely chemotaxis and migration (**Fig. 2H**). There were particularly marked changes in the expression of genes such as CBLN2 which can promote EndMT by activating the NF-κB/HIF-1α/Twist1 pathway,^14^ and NEXN (**Fig. 2I**), which encodes nexilin, an actin binding protein which like SMOC1 has been implicated in bone differentiation^15^ and regulates vascular smooth muscle cell phenotypic switching and neointimal hyperplasia.^16^

SMOC1 belongs to the SPARC (Secreted Protein Acidic and Rich in Cysteine) family of matricellular proteins that can dynamically modulate cell-ECM interactions. SMOC1 can bind matrix proteins such as tenascin C^17,18^ and can interfere with extracellular BNP and TGF-βsignaling.^13,19–21^ Therefore, we determined whether it was possible to reverse the effects of SMOC1 downregulation using a recombinant SMOC1 protein (rSMOC1). The deletion of SMOC1 aggravated the EndMT phenotype, potentiating the cytokine-induced decrease in CD144 and increase in SM22α RNA (**Fig. 3A**) and protein, confirmed by increased number of SMA^+^ endothelial cells (**Fig 3B**). Both effects were abrogated by the addition of rSMOC1, as were EndMT-associated changes in cell size (**Fig. 3C**).

**Fig. 3.**
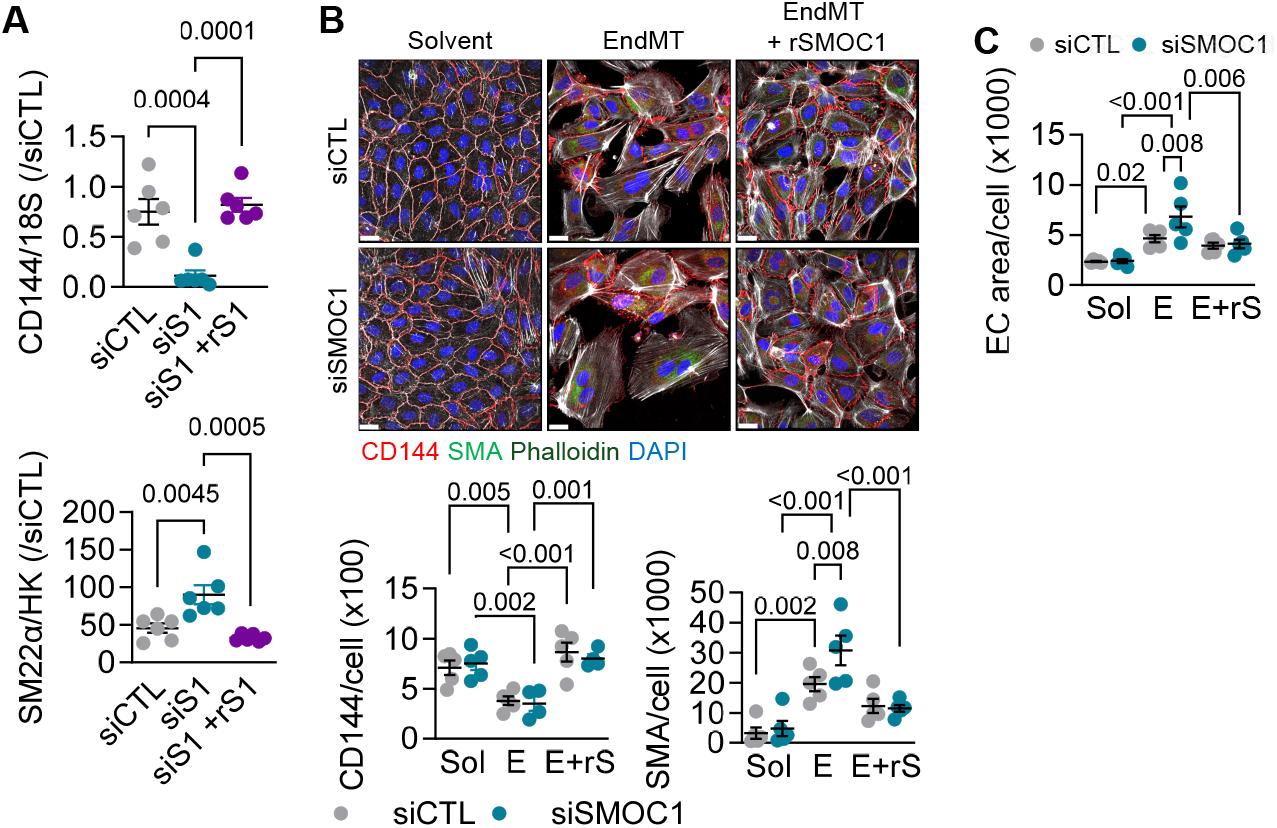
Recombinant SMOC1 rescues the effects of SMOC1 downregulation on EndMT induction. (**A&B**) Impact of SMOC1 downregulation (siS1) versus control (siCTL) on CD144 and SM22α or αSMA RNA (A) and protein (B) expression in endothelial cells in EndMT. Experiments were performed in the absence and presence of rSMOC1; n=5-6 independent cell batches. A: One way ANOVA and Tukey’s multiple comparisons test. B: Two way ANOVA and Tukey’s multiple comparisons test. (**C**) Impact of SMOC1 downregulation (siS1) versus control (siCTL) on the size of cells in EndMT and rescue by rSMOC1 on cell size; n=6 independent cell batches. Two way ANOVA and Tukey’s multiple comparisons test.

As rSMOC effectively rescued EndMT, we assessed its ability to interfere with TGF-β2 signaling. Consistent with the previously reported ability of SMOC1 to redirect TGF-β1 signaling from ALK5-SMAD2 to ALK1-SMAD1, the TGF-β2-induced phosphorylation of SMAD2 was clearly decreased in SMOC-1 deficient cells (**Fig. 4A**). In the same cells there was a subtle but significant increase in the phosphorylation of SMAD1/5. The addition of rSMOC1 reversed the effects of SMOC1 deletion and attenuated the ability of TGF-β2 to elicit the phosphorylation of SMAD2 (**Fig. 4B**). SMOC1 was previously found to alter ALK5 signaling by binding to endoglin (CD105),^13^ the expression of which was also increase by EndMT (see Fig 2A&B). Membrane-bound endoglin can exist as a long and short form and we found that EndMT induced a highly significant increase in the short isoform (S-endoglin) without significantly altering long (L)-endoglin (**Fig. 4C**). This is of relevance as L-endoglin is thought to be the signaling competent isoform and that S-endoglin, by virtue of its truncated cytoplasmic tail, lacks relevant signaling domains^22,23^ and responds differently to TGF-β.^24,25^ Adding rSMOC1 to the EndMT induction cocktail switched the isoforms, resulting in decreased S-endoglin and increased L-endoglin.

**Fig. 4.**
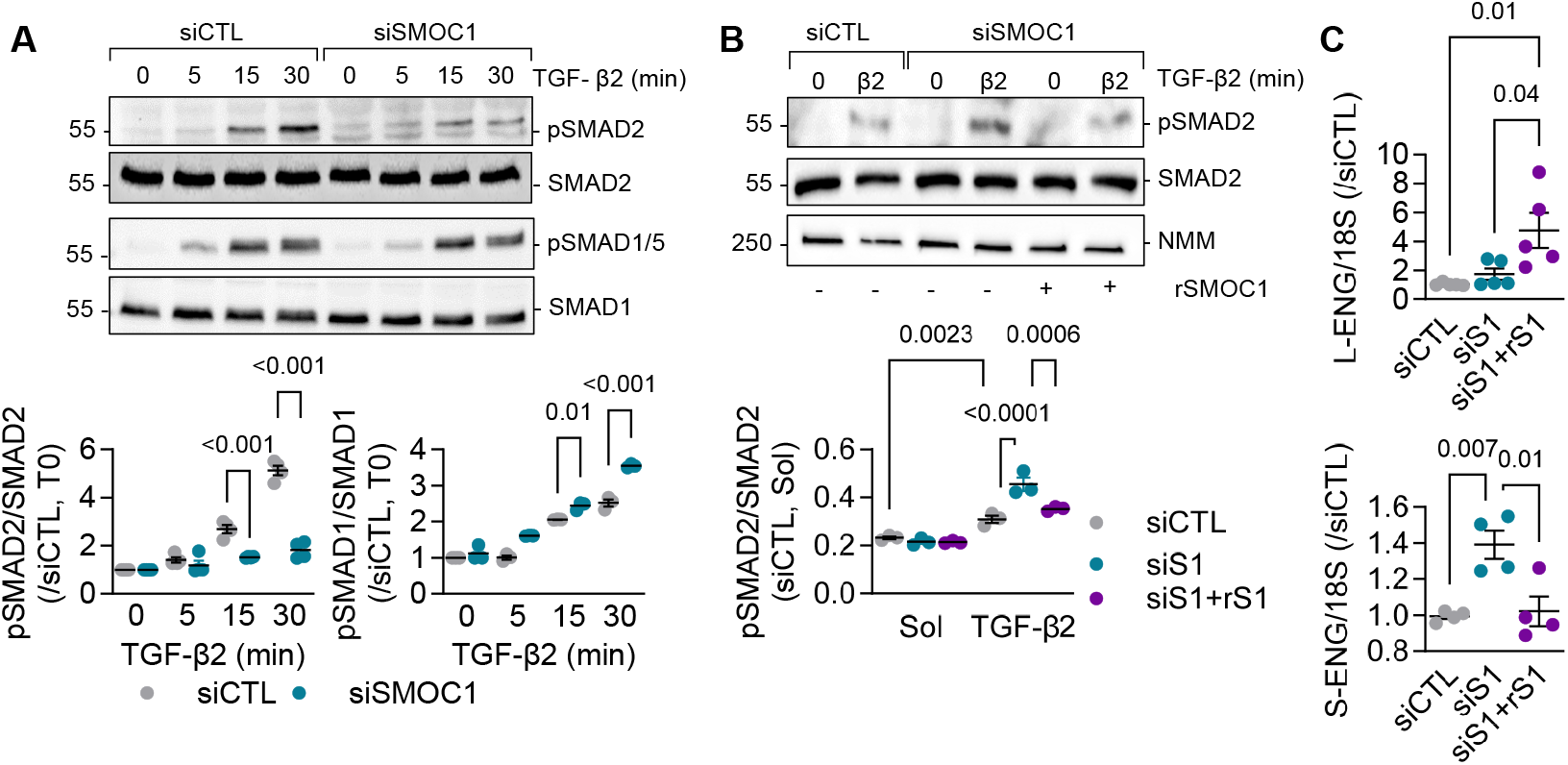
Impact of SMOC1 downregulation and rescue with rSMOC1 on TGF-β2 signaling. (**A**) Impact of SMOC1 downregulation (siSMOC1) on the TGF-β2-induced phosphorylation of SMAD2 and SMAD1/5; n=3 independent cell batches. Two way ANOVA and Sidak’s multiple comparisons test. (**B**) Effect of rSMOC1 (rS1) on the TGF-β2-induced phosphorylation of SMAD2 in SMOC1-deficient cells (siSMOC1); n=3 independent cell batches. Two way ANOVA and Tukey’s multiple comparisons test. (**C**) Expression of L-endoglin (L-END) and S-END in cells treated with a control siRNA (siCTL) or siRNA directed against SMOC1 (siS1) in the absence and presence of rSMOC1 (rS1); n=5 independent cell batches. One way ANOVA and Bonferroni’s multiple comparisons test.

Several studies have made a connection between aging and EndMT as they share initiating factors such as oxidative stress and inflammation (for review see reference^26^). Therefore, we determined whether the deletion of SMOC1 could precipitate EndMT in aged mice. Because SMOC1^-/-^ mice do not survive for long after birth^27–30^ and SMOC1^+/-^ mice express little or no detectable protein,^13,31^ we maintained wild-type and SMOC1^+/-^ littermates on a standard chow diet for 1 year and then assessed SMAD2 phosphorylation in aortic endothelial cells as an indicator of EndMT. This revealed that SMOC1-deficiency increased SMAD2 phosphorylation, particularly in the small curvature of the aorta (**Fig. 5A**). Because SMOC1 can be generated by other cell types e.g. platelets,^31^ we next generated mice lacking SMOC1 specifically in endothelial cells (SMOC1^ΔEC^ mice). Also in these animals there was evidence of spontaneous aging-associated EndMT in the form of the expression of SM22α (**Fig. 5B**). RNA-sequencing of endothelial cells from these mice confirmed the induction of an EndMT state as evidenced by the increased expression of TGF-β3, Snai2 and others (**Fig. 5C**).

**Fig. 5.**
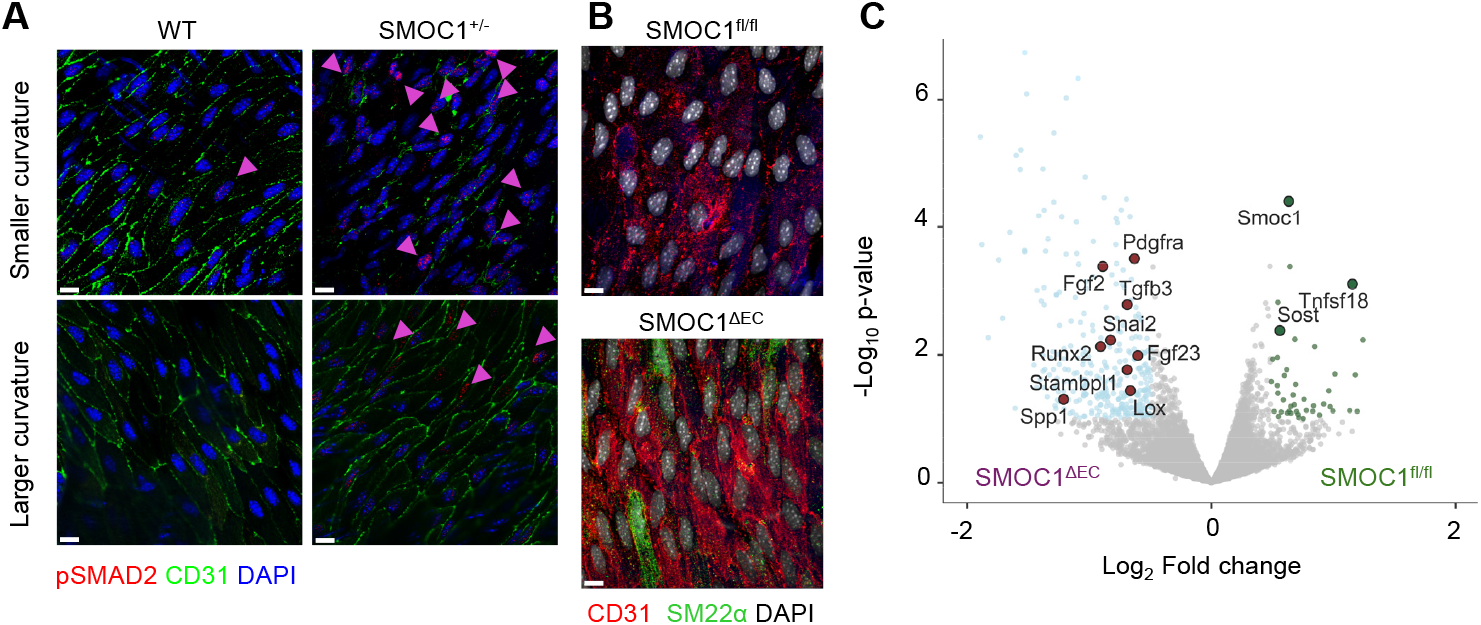
EndMT in wild-type and SMOC1-deficient mice. (**A**) En face images of the small and large curvatures of the aortic arch from 1 year old SMOC1^+/+^ and SMOC1^+/-^ littermates. Comparable results were obtained in an additional 3 mice per genotype. Arrowheads highlight the positive pSMAD2 signal. Bar = 10 µm. (**B**) En face images of the small and large curvatures of the aortic arch from 1 year old wild-type (SMOC1^fl/fl^) mice and mice lacking SMOC1 specifically in endothelial cells (SMOC1^ΔEC^). Comparable results were obtained in an additional 3 mice per genotype. Bar = 10 µm. (**C**) Volcano plot showing DEGs in endothelial cells isolated from 1 year old SMOC1^fl/fl^ and SMOC1^ΔEC^ mice; n=6 mice per genotype.

Given the contribution of EndMT in both initial scar stabilization and subsequent pathological fibrosis, we assessed the consequence of deleting endothelial cell SMOC1 on the response to acute MI. After infarction, SMOC1^fl/fl^ and SMOC1^ΔEC^ mice experienced a decline in systolic function and an increase in left ventricular end diastolic volume (**Fig. 6A**). However, these changes were significantly aggravated by SMOC1 deletion. Fourteen days after MI, SMOC1^ΔEC^ mice exhibited marked left ventricular dilation and wall thinning (**Fig. 6B**), with a significant loss of viable myocardium and an increased collagen-rich fibrotic area (**Fig. 6C**). In several instances, cardiac rupture occurred and survival was reduced in the SMOC1ΔEC group (**Fig. 6D**). Because endothelial cells orchestrate inflammatory signaling during post-ischemic repair, we next assessed immune cell infiltration during the acute phase. Flow cytometric profiling 1 and 3 days after MI revealed an increase in multiple inflammatory cell subsets, including neutrophils, monocytes, macrophages and dendritic cells in hearts from SMOC1^ΔEC^ versus SMOC1^fl/fl^ mice (**Fig. 6E**).

**Fig. 6.**
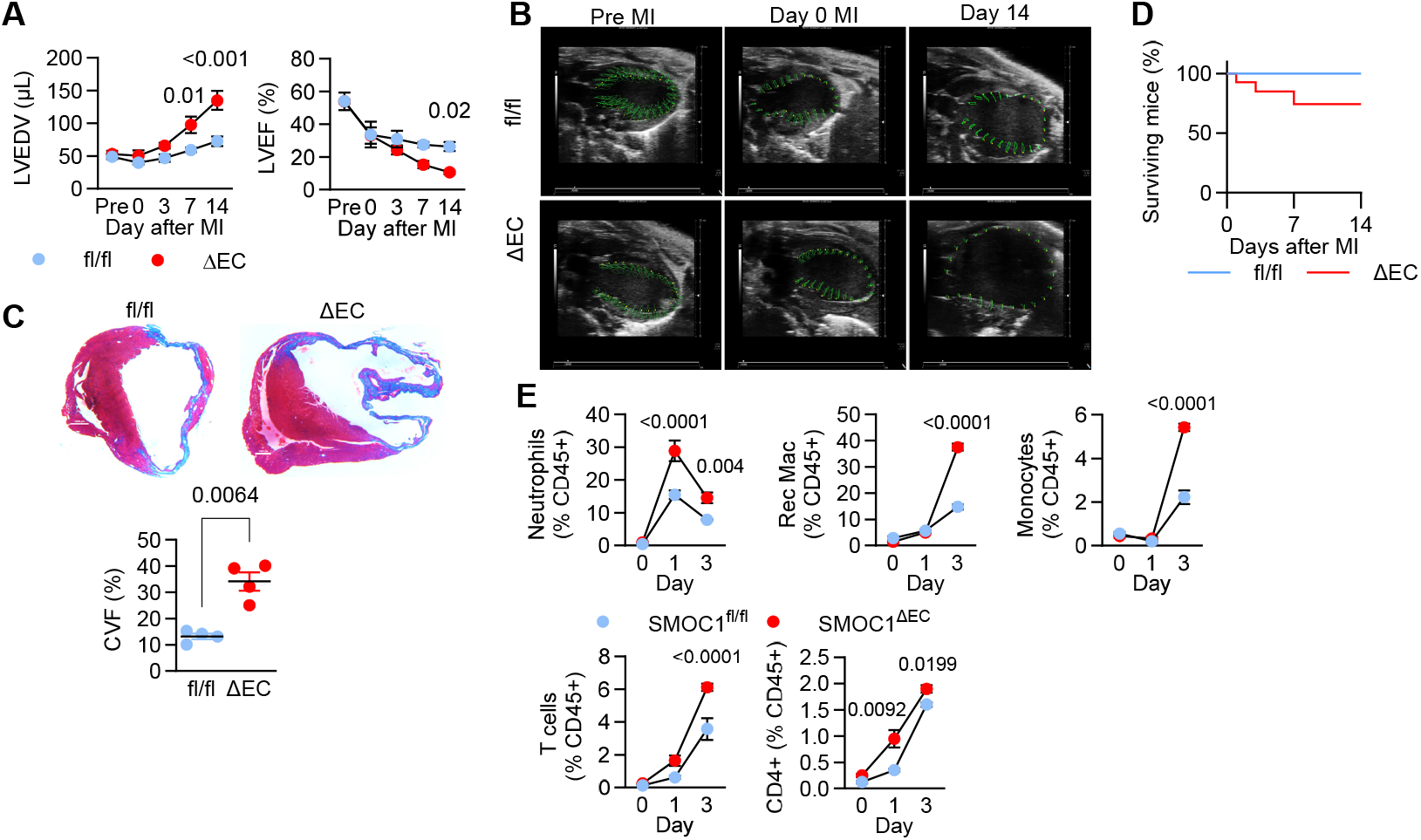
Impact of endothelial cell-specific SMOC1 deletion on the response to MI. (**A**) Changes in left ventricular end diastolic volume (LVEDV) and left ventricular ejection fraction (LVEF) in SMOC1^fl/fl^ and SMOC1^ΔEC^ at baseline (pre) and up to 14 days after MI; n=8 mice per group. Two way ANOVA with Bonferroni’s multiple comparisons test. (**B**) Representative M-mode echocardiographic images from SMOC1^fl/fl^ (fl/fl) and SMOC1^ΔEC^ (ΔEC) mice. Overlaid green vectors indicate the movement (strain/velocity) of the cardiac muscle during the cardiac cycle. (**C**) Masson’s Trichrome staining of hearts from SMOC1^fl/fl^ and SMOC1^ΔEC^ mice 14 days post-MI; n=4 mice / genotype. Student’s t test. (**D**) Kaplan-Meier plot showing survival of the 2 strains after MI; n = 10 mice per genotype. (**E**) Immune cell infiltrations into hearts up to 3 days after MI; n= 4 mice / genotype. Two way ANOVA with Bonferroni’s multiple comparisons test.

## Discussion

The results of the present investigation identify SMOC1 as a key endothelial cell-derived regulator of EndMT *in vitro* and *in vivo*, with profound consequences for post-infarct cardiac remodeling and function. The lack of endothelial cell-derived SMOC1 had a negative impact on the myocardial response to injury, with significantly greater fibrosis and incidence of cardiac rupture. At the molecular level, SMOC1 interfered with TGF-β2 to prevent the agonist-induced phosphorylation of SMAD2. Consistent with an extracellular action of SMOC1 these effects could be reversed by the addition of rSMOC1. Intriguingly, in the absence of SMOC1, the predominant endoglin isoform switched from the long form (L-endoglin) to the short form (S-endoglin). While S-endoglin has a truncated cytoplasmic tail and altered signaling properties, our data indicate that its predominance in the absence of SMOC1 is associated with a net shift toward enhanced pro-fibrotic TGF-β/SMAD2/3 signaling. This suggests that SMOC1 may fine-tune the cellular response to TGF-β2 by modulating the endoglin receptor complex, thereby diverting signaling away from a profibrotic, pro-EndMT SMAD2/3 axis.

SMOC1 has recently gained a lot of attention as it is one of the most highly upregulated proteins in brain, cerebrospinal fluid and plasma samples from patients with Alzheimer’s disease,^32–35^ and has been proposed to play a role in the regulation of endothelial and blood hemostatic pathways linked to blood brain barrier disruption.^36^ However, whether it plays a functional role in disease development is not clear. Not a lot is known about the role of SMOC1 in the cardiovascular system, but it plays a role in development, including cardiac development.^18,27,37^ There are, however, a few reports of altered SMOC1 levels associated with pathologies. For example, SMOC1 was expressed by aortic valve endothelial cells and secreted into the ECM of non-calcific valves and downregulated in calcific aortic valves.^19^ Platelets can also express the protein and an increase in its production was proposed to underlie enhanced reactivity to thrombin observed in platelets from diabetic individuals and mice.^31^ In fact, SMOC1 is one of the few proteins that can bind to and activate thrombin.^31^ Moreover, SMOC1 expression is reportedly elevated in subjects with hypertension,^38^ for which it has been proposed as a useful biomarker^39^.

SMOC1 is a secreted protein that binds to numerous components in the ECM as well as to growth factors such as bone morphogenic proteins and TGF. It can also bind to membrane proteins to alter responses to the same agonists^13,20,40–42^ and to-date SMOC1 was found to associate with the BMP receptor II^19^ as well as the auxiliary TGF-β receptor endoglin.^13^ There may however be other consequences of SMOC1 gene expression that are independent of the secreted protein as the SMOC1 sequence hosts a competitive endogenous RNA (miR-211-5p-SMOC1) that was proposed to play a crucial role in the occurrence and development of coronary artery disease,^43^ and a circular RNA reported to regulate vascular smooth muscle calcification^44^ and to promote pulmonary hypertension.^45^

In the present study we identified a robust upregulation of SMOC1 in a population of cardiac EndMT cells following MI, a phenomenon that correlated temporally with the upregulation of well-defined EndMT marker genes. Given the central role of TGF-β signaling in the EndMT process (for review see^26,46^) and the fact that SMOC1 is a TGF-β regulated gene,^42^ this was perhaps to be expected. However, we observed that SMOC1 played a functional role and tempered the mesenchymal transition of endothelial cells as its downregulation resulted in an exaggerated response and elevated induction of EndMT marker genes like CBLN2 and NEXN. Importantly, all of these effects were attributable to the SMOC1 protein rather than any of the associated RNA species as the consequences of SMOC1 knockdown on EndMT could be prevented by the recombinant protein.

The events linking extracellular SMOC1 and intracellular changes in gene expression seem to be determined by its impact on BMP and TGF-β signaling. Indeed, it was previously found to bind to endoglin to antagonize TGF-β1-ALK5-SMAD2 signaling and activate that of ALK1-SMAD1/5 to stimulate angiogenesis.^13^ Our observations using IL-1β and TGF-β2 to induce EndMT confirm the latter report and are suggestive of a similar switch in TGF-β signaling. However, we also observed that endoglin levels increased in parallel with those of SMOC1 and that an isoform switch from L-to S-endoglin occurred. While L-endoglin is thought to supports vascular health by promoting nitric oxide synthase (eNOS) expression and stabilizing TGF-β signaling, S-endoglin tends to inhibit these pathways, contributing to vascular dysfunction and disease progression.^47,48^ There is circumstantial evidence linking endoglin with EndMT^49–51^ but there has been no detailed analysis of the role of the different isoforms. Given that the S-endoglin that was elevated in endothelial cells in EndMT carries a truncated cytoplasmic tail, it is tempting to suggest that the changes in both proteins play a fundamental role in EndMT.

The process of EndMT plays a crucial role in the response to damage and its cessation or reversion is a key component of the repair process and the restoration of a homeostatic state.^4^ Overall, our study indicates that SMOC1 serves a protective homeostatic function, restraining EndMT activation under chronic pro-aging stimuli like inflammation and oxidative stress. Indeed, even in the absence of injury, the lack of endothelial cell-derived SMOC1 could be associated with spontaneous EndMT of aortic endothelial cells from aged SMOC1^ΔEC^ mice, which expressed SM22α and displayed basal SMAD2 phosphorylation, particularly in hemodynamically stressed regions of the aorta. Following MI, this protective role became critical as SMOC1^ΔEC^ mice suffered worse outcomes with exacerbated ventricular dilation, wall thinning, fibrosis, and increased mortality due to cardiac rupture. This accelerated maladaptive remodeling was coupled with a significantly amplified acute inflammatory response, with heightened infiltration of neutrophils, monocytes, and macrophages. This suggests that SMOC1 in endothelial cells modulates not only cell-autonomous mesenchymal-like transition but also the endothelial control of post-ischemic inflammation, two processes that are intimately linked in driving adverse remodeling.

## Acknowledgements

The authors are indebted to Isabel Winter, Mechtild Piepenbrock-Gyamfi and Katharina Bruch for expert technical assistance. This work was supported by the Deutsche Forschungsgemeinschaft (SFB1531/1 B03, A05, A03 – Project ID 456687919; GRK 2336 TP5

-Project ID 321115009; CardioPulmonary Institute, EXC 2026, Project ID: 390649896) and the Dr. Rolf M. Schwiete Stiftung (Projekt Nr. 2021-025).

## Materials and methods

### Animals

C57BL/6J wild-type mice (>8 weeks, >18 g) were obtained from Charles River (Sulzfeld, Germany). SMOC1^fl/fl^ mice (Smoc1tm1a(EUCOMM)Wtsi) were acquired from TaconicArtemis (Cologne, Germany) and crossed with C57BL/6-Tg(ACTB-Flpe)2Arte animals to remove the FRT-flanked LacZ/neo cassette. Endothelial-specific SMOC1 knockout mice (SMOC1^ΔEC^) were generated by breeding SMOC1^fl/fl^ mice with B6.FVB-Tg(Cdh5-cre)7Mlia/J mice from the Jackson Laboratory (Bar Harbor, ME, USA). Mice with heterozygous knockout of the SMOC1 gene, i.e., expressing severely reduced levels of SMOC1, were obtained from the RIKEN BioResource Centre (Tsukuba, Japan). Animals were housed under a 12-hour light-dark cycle with free access to water and food. All animal experiments were performed in accordance with the Directive 2010/63/EU of the European Parliament on the protection of animals used for scientific purposes and approved by the Federal Authority for Animal Research at the Regierungspräsidium Darmstadt (Hessen, Germany) under study protocol FU/2047. Age-, sex-, and strain-matched animals (littermates) were used throughout the studies. For the isolation of heart, mice were euthanized using 4% isoflurane in air and subsequently exsanguinated.

### Myocardial infarction

Myocardial infarction (MI) was induced by minimally invasive procedure using echocardiographic guidance described previously.^52^. Briefly, mice were anesthetized with 2% isofluorane and placed on the warming pad of Vevo 3100 ultrasound imaging system (VisualSonics, Toronto, Canada) to maintain 37°C body temperature. Baseline cardiac function and morphology were evaluated before MI induction. After color doppler visualization, the left anterior descending coronary artery was electrically cauterized with a 26G monopolar needle (#027722, Natus Europe GmbH, Planegg, Germany) inserted in the chest using a micromanipulator. Successful coronary artery occlusion and MI were confirmed by absence of color doppler signal and akinesia in the left ventricular wall. Mice were laid in a prone position on warming pad and then transfered to a new cage after awakening.

### Echocardiography

Cardiac function and morphology were assessed on day 0, 3, 7, and 14 after MI by two-dimensional echocardiography using Vevo 3100 ultrasound imaging system (Vevo software version 3.2.8) equipped with MS550D 22-50 MHz linear array solid-state transducer (VisualSonics, Toronto, Canada). Mice were anesthetized with 2% isoflurane, maintaining body temperature and heart rate at 37°C and over 450 bpm for all the time of the measurement. Two-dimensional B-mode echocardiography was performed in parasternal long- and short-axis views, and images were analyzed using a modified Simpson approach. From these multi-planar tracings, left ventricular anterior and posterior wall thickness, septal thickness, and left ventricular internal diameters at end-systole and end-diastole were obtained, and these measurements were then used to calculate left ventricular fractional shortening and ejection fraction using Vevo LAB 5.6.0 software (Visualsonics, Toronto, Canada), according to ESC guidelines.^53^

### Flow cytometry profiling of cardiac immune cells

One and 3 days after MI hearts were harvested, residual connective tissue removed and cells were dissociated using a MACS Octo Dissociator (Miltenyi Biotec, Bergisch Gladbach, Germany), filtered through a 70-µm cell strainer, centrifuged and sequentially processed with Miltenyi Debris Removal kit (Cat. #: 130-109-398, Miltenyi Biotec, Bergisch Gladbach, Germany) and Red Cell Lysis kit (Cat #:130-094-183 Alfa Aesa, Kandel, Germany). The final pellet was resuspended in flow cytometry buffer (2% bovine serum albumin (BSA), 1 mmol/L ethylenediaminetetraacetic acid (EDTA) in phosphate-buffered saline (PBS)). For staining, 100 µl of the cell suspension was washed in PBS/0.5% BSA, blocked with 1:200 FcR-Binding-Inhibitor(Cat. #: 130-092-575, Milteny, Bergisch Gladbach, Germany), and incubated with an antibody master mix diluted 1:500 PBS/0.5% BSA including: CD11b-Alexa Fluor 647 (rat, Cat. #: 101220), CD11b-BV421 (rat, Cat. #: 101251), CD11b-BV605 (rat, Cat. #: 100742), CD192/CCR2-PE (rat, #150610), F4/80-PE-Cy7 (rat, #123114), Ly6G-APC-Cy7 (rat, Cat. #: 127624), and Ly6G-PE (rat, Cat. #: 127607), all from BioLegend (Koblenz, Germany); Ly6C-PerCP-Cy5.5 (rat, Cat. #: 560525) and CD3-PE-CF594 (Armenian hamster, Cat. #: 562286) from BD Biosciences (Heidelberg, Germany); and CD45-VioBlue (rat, Cat. #: 130-102-430) from Miltenyi Biotec (Bergisch Gladbach, Germany). After incubation at 4 °C for 25 min in the dark, cells were washed, resuspended in flow cytometry buffer with counting beads, and analysed within 3 hours on an LSRII/Fortessa cytometer (BD Bioscience, Heidelberg, Germany) and the data was analyzed in FlowJo™ v10.9.0 Software (BD Life Sciences, Heidelberg, Germany).

### Tissue collection and histological analysis

#### Sample preparation

At day 14 post-MI, mice were anesthetized with 4% isoflurane and euthanized. Hearts were collected, washed with PBS, fixed overnight with 4% paraformaldehyde at 4 °C and processed for paraffin embedding for histological analysis. Using 5 µm-thick sections, tissue fibrosis was assessed by Masson-Goldner trichrome staining (Cat. #: HT15-1KT, Merck, Darmstadt, Germany), following the manufacturer’s protocol. All histological sections were digitized using a microscope equipped with a high-resolution camera (AxioObserver.Z1, Zeiss). To ensure comparability, identical exposure settings and a constant 4× magnification were used for all samples. Images were subsequently processed and archived using Zen Blue 6.0.3 (Zeiss).

#### Histological analysis

Scar size was assessed on five short-axis series per heart, evenly spaced between the most basal section showing the complete left ventricle (LV) with papillary muscles and the most apical section in which the LV was still fully visible. At each level, one Masson-Goldner-stained section was analyzed in ImageJ (NIH). Total LV myocardium and infarcted (fibrotic) tissue were delineated by color thresholding, and pixel areas were recorded. For each section, infarct percentage was calculated as scar area divided by total LV myocardial area. Final infarct size per animal was obtained as the mean infarct percentage of the five sections multiplied by the ratio of infarcted to total series, considering a level infarcted when scar tissue was detectable microscopically.

### RNA sequencing

#### Bulk RNA-seq

After a three-day EndMT induction period, endothelial cells were incubated with a lysis buffer containing 1% NP-40, 50 mmol/L Tris-HCl (pH 7.5), 150 mmol/L NaCl, 10 mmol/L MgCl_2_ phosphatase and protease inhibitors (10 mmol/L NaPPi, 20 mmol/L NaF, 10 nmol/L okadaic acid, 2 mmol/L Na_3_VO_4_, 12 µl/mL PIM, 4 µl/mL PMSF), 200 U/mL SUPERase•In RNase inhibitor and 25 U/mL TURBO DNase I in UltraPure DNase/RNase-free water for 45–60 minutes at 4°C. Complete membrane disruption was achieved by mechanically shearing the suspension through a 26G needle attached to a 1-mL syringe for 7–10 passes. The lysate was clarified by centrifugation at 13,000 rpm for 10 minutes at 4°C, and the resulting supernatant was used for total RNA isolation with the miRNeasy Micro Kit according to the manufacturer’s instructions.

RNA integrity and library preparation quality were evaluated using a LabChip Gx Touch system. For library generation, 500 ng to 1 µg of total RNA was processed using the SMARTer Stranded Total RNA Sample Prep Kit HI Mammalian. Sequencing was conducted on an Illumina NextSeq 2000 using a P3 flowcell, producing 72-bp single-end reads.

Preprocessing of raw reads was performed with Trimmomatic (v0.39) to remove low-quality regions (sliding-window trimming at Q15 over 5 nucleotides) and to retain sequences ≥15 nt. Filtered reads were aligned to the human reference genome (Ensembl hg38, release 109) using STAR (v2.7.10a). Post-alignment cleanup involved removing duplicate, multi-mapping, ribosomal, and mitochondrial reads with Picard (v3.0.0). Gene-level quantification was carried out using featureCounts (v2.0.4) with exon-based, strand-specific read assignment. Differential expression analysis was conducted in DESeq2 (v1.36.0) on untransformed counts. Genes were defined as significantly regulated when exhibiting an average count >5, an adjusted p-value <0.05, and an absolute log2 fold change ≥0.585. Gene annotations from Ensembl were complemented with UniProt metadata.

#### Bulk RNA-seq data analysis

Functional enrichment was performed using clusterProfiler (v4.16.0) with the GO:BP database (org.Hs.eg.db v3.21.0). Over-Representation Analysis (ORA) of DEGs was performed using enrichGO (BH-adjusted p < 0.05, q < 0.2). Gene Set Enrichment Analysis (GSEA) was conducted on the full gene list ranked by averaged Wald statistic using gseGO (set size 10–500, BH-adjusted p<0.05). Results were visualized using enrichplot. Global expression was visualized via PCA on VST-normalized counts using 95% confidence ellipses. Targeted heatmaps were generated with ComplexHeatmap, displaying Z-score scaled expression for defined gene signatures and the TGF-beta response pathway, with samples ordered by condition. Marker expression (FN1, SMOC1, TAGLN, PECAM1) was visualized using violin plots, annotated with DESeq2 Wald test statistics (BH-adjusted p-values)

#### Single-cell RNA-seq data analysis

Public datasets with the following GEO accession numbers were used: GSE176092, GSM5355657, GSE207289 and GSE216211. Single-cell RNA sequencing data processing and analysis were performed using R (v4.5) and the Seurat package. Quality control steps included filtering cells based on mitochondrial content and detected feature counts. Data were log-normalized, scaled, and highly variable features were identified for dimensionality reduction via Principal Component Analysis (PCA). Non-linear visualization was performed using Uniform Manifold Approximation and Projection (UMAP), followed by graph-based clustering.

Endothelial cells were computationally subsetted for high-resolution sub-clustering. Pathway activity scores for specific signatures, including TGF-beta signaling and EndMT, were calculated at the single-cell level using the AddModuleScore function. Additionally, functional pathway enrichment was assessed using Fast Gene Set Enrichment Analysis (FGSEA) against the MSigDB Hallmark collection to identify statistically significant biological processes associated with specific timepoints and conditions.

### Cell culture

#### Human endothelial cells

Human umbilical cords were collected from local hospitals and umbilical vein endothelial cells were isolated and cultured following previously established procedures^54,55^ and were used only up to passage 4. All work involving human-derived material adhered to the ethical standards of the Declaration of Helsinki,^56^ and the collection and use of endothelial cells were approved in writing by the Ethics Committee of Goethe University. Endothelial cells were cultured in ECGM2 medium (PromoCell, Heidelberg, Germany) containing 8% heat inactivated foetal bovine serum (FBS), gentamycin (25 µg/mL) non-essential amino acids (Thermo Fisher Scientific, Schwerte, Germany) and Na pyruvate (1 mmol/L, Merck, Darmstadt, Germany).

#### EndMT induction

Human endothelial cells were treated with a combination of IL1-β (1 ng/mL, Cat #: 200-01B Peprotech, Hamburg, Germany) and TGF-β2 (1ng/mL, Cat. #:100-35B Peprotech, Hamburg, Germany) for 72 hours, replacing the culture medium daily to induce EndMT. For rescue experiments, recombinant SMOC1 (rSMOC1, 0.5µg/mL, Cat. #:6074-SM-050, R&D system, Wiesbaden, Germany) was added 30 minutes prior to the TGF-β2.

#### SMOC1 knockdown

Endothelial cells were transfected with 50 nM small interfering RNA (siRNA) targeting SMOC1 (sense: 5′-UUG UUA AUG UCG UUG CUG C-dTdT-3′; antisense: 5′-GCA GCA ACG ACA UUA ACA UUA ACA A-dTdT-3′) or with corresponding control oligonucleotides (Eurogentec, Cologne, Germany). All transfections were carried out using the Lipofectamine RNAiMax Transfection Kit (Invitrogen, Darmstadt, Germany) according to the manufacturer’s protocol.

### Immunofluorescence

Endothelial cells were washed with PBS, fixed in 4% paraformaldehyde (Carl Roth) at room temperature for 15 minutes, permeabilized with 0.1% Triton X-100 and subsequently blocked with 3% BSA in PBS (Roth, Cat. #:8076.3) for 1h at room temperature. Cells were then incubated with primary antibodies diluted in 3% BSA overnight at 4 °C. Goat- and mouse-specific secondary antibodies diluted (1:200) in PBS were applied for 1 hour and nuclei were stained with DAPI (10 ng/mL), before mounting using Fluoromount-G (Cat. #: 00-4958-02; Invitrogen, Thermo Fisher Scientific, Waltham, MA, USA). Images were taken with a confocal microscope (Leica SP8 or Zeiss LSM-780) and analyzed using the respective software i.e., LASX (Server version 1.9.0.3233, Leica, Wetzlar, Germany) or ZEN 2012 (blue edition) software (Zeiss, Jena, Germany) as described.^57^

### Cell lysis and immunoblotting

Proteins were extracted using RIPA lysis buffer (50 mmol/L Tris/HCl, pH 7.5, 150 mmol/L NaCl, 10 mmol/L NaPi, 20 mmol/L NaF, 1% sodium deoxycholate, 1% Triton X-100, and 0.1% SDS) supplemented with protease and phosphatase inhibitors. Briefly, cells were lysed in ice-cold lysis buffer containing 50 mmol/L Tris/HCl (pH 7.5), 150 mmol/L NaCl, 1% NP-40, 10 mmol/L NaPPi, 20 mmol/L NaF, 2 mmol/L sodium orthovanadate, 10 nmol/L okadaic acid, 25 mmol/L β-glycerophosphate, and 230 µmol/L PMSF. Lysates were centrifuged at maximum speed (≈15,000 rpm) for 15 min at 4 °C, and the supernatant was collected and stored for further analysis. Detergent-soluble proteins were resuspended in SDS-PAGE sample buffer, separated using SDS-PAGE, and processed for Western blotting as previously described.^54^ using the antibodies listed in Table 2. Membranes were incubated with species-specific horseradish peroxidase–conjugated secondary antibodies diluted 1:20,000 in Tris-buffered saline–Tween. Protein bands were then visualized by enhanced chemiluminescence using a commercial ECL kit (GE Healthcare, Chicago, United States).

**Table 1.**
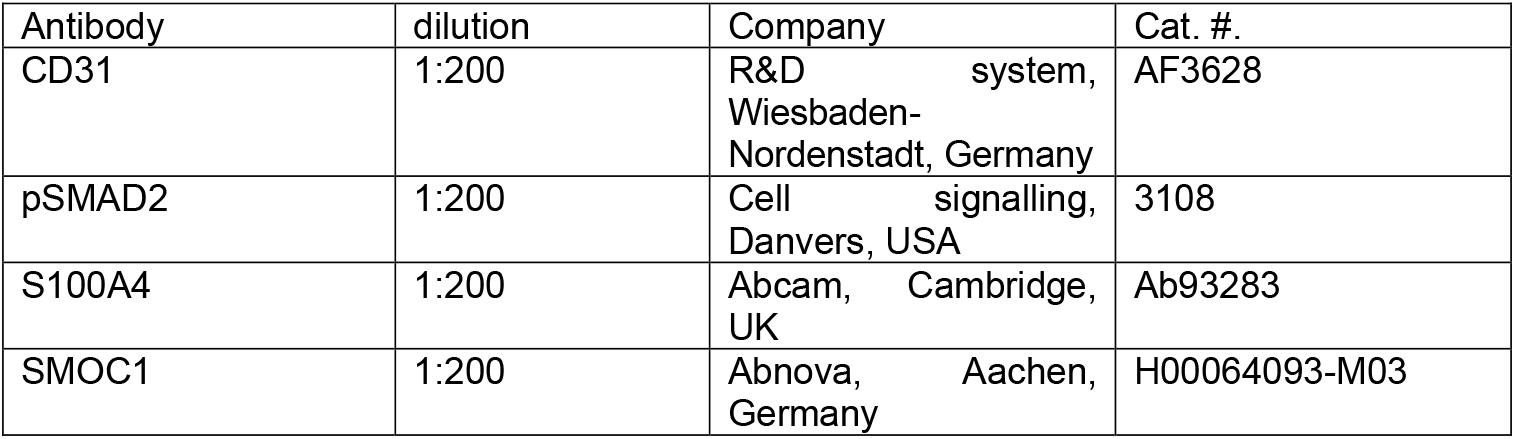
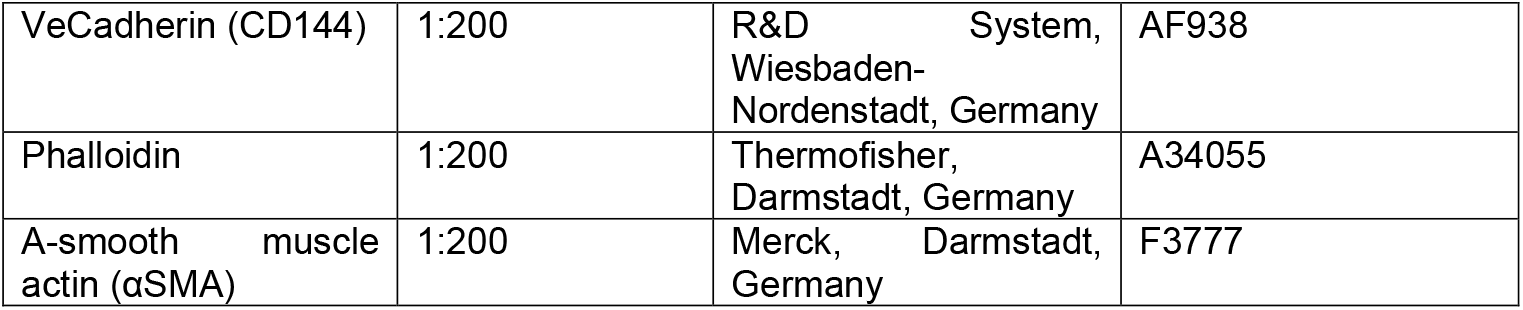
Antibodies used for immunofluorescence.

**Table 2.** Antibodies used for Western blotting.

| Antibody | dilution | Company | Cat No. |
| --- | --- | --- | --- |
| pSMAD2 | 1:1000 | Cell signalling,<br>Danvers, USA | 3108 |
| pSMAD1/5 | 1:1000 | Cell signalling,<br>Danvers, USA | 9516 |
| SMAD2 | 1:1000 | Cell signalling,<br>Danvers, USA | 5339 |
| SMAD1 | 1:1000 | Cell signalling,<br>Danvers, USA | 6944 |
| NMM | 1:1000 | DPC Biermann | BP_643 |
| SMOC1 | 1:1000 | Abnova, Aachen,<br>Germany | H00064093-M03 |
| B-Actin | 1:1000 | Sigma, Munich,<br>Germany | A1978 |

**Table 3.**
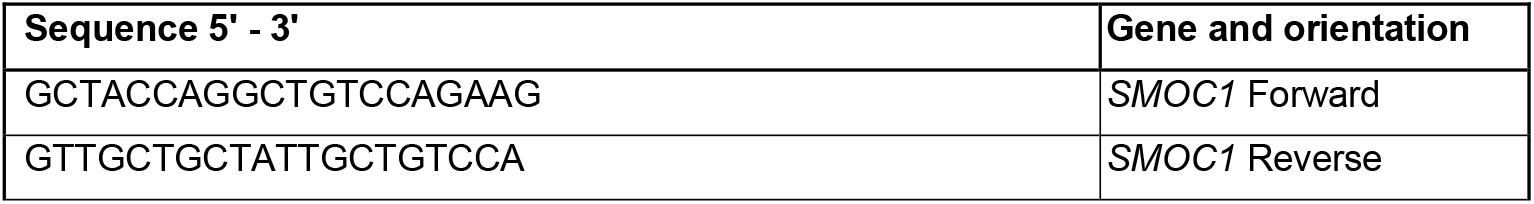
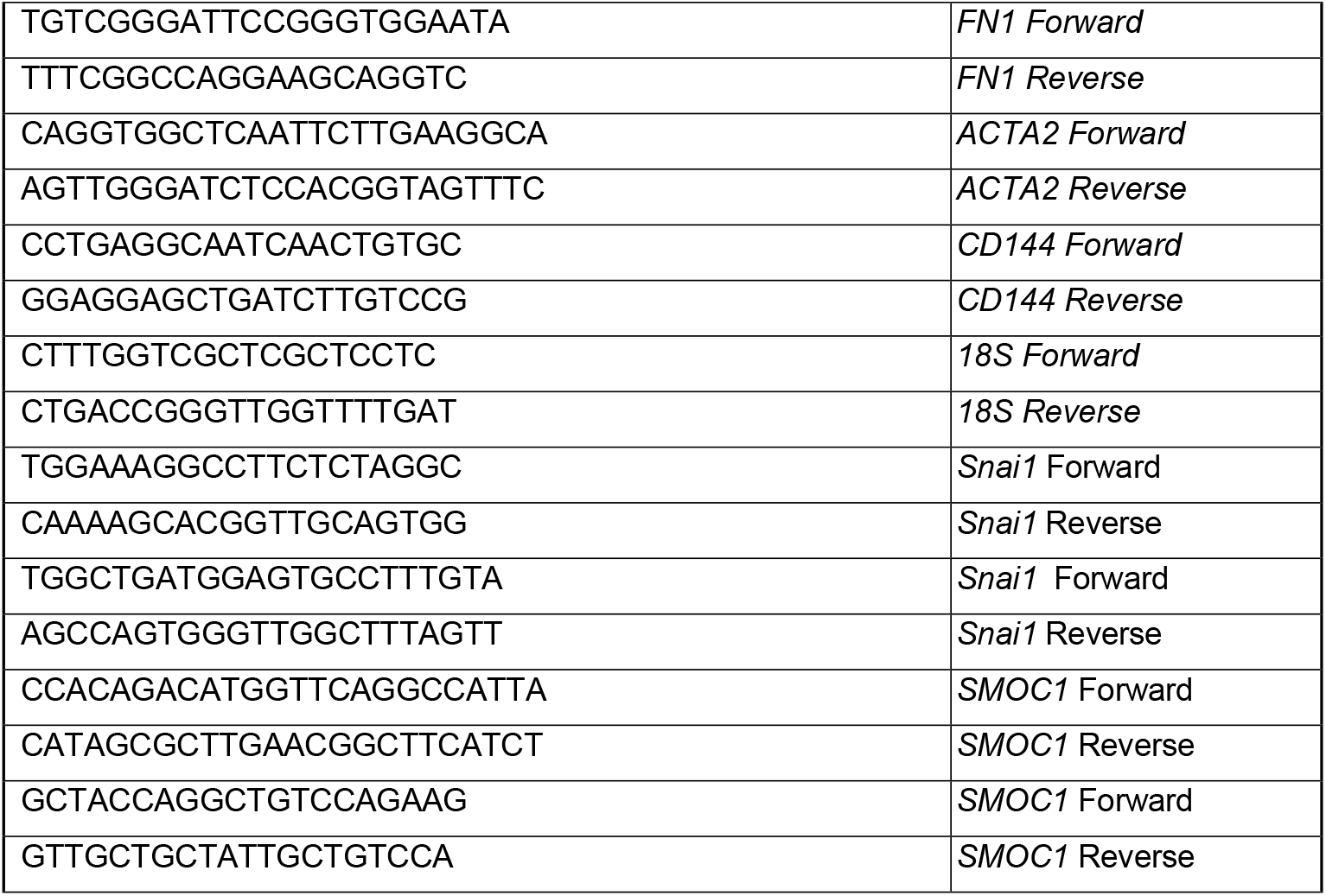
Oligonucleotides used for RT-qPCR.

### RT-qPCR

Total RNA was extracted using TRIzol (Invitrogen, Cat. #: 15596018) and retrotranscribed to cDNA using First Strand cDNA Synthesis Kit (Thermo Scientific, Cat. #: K1612) with random hexameric primers following the manufacturer’s instructions. RT-qPCR was performed using SYBR green master mix (Biozym, Hessisch Oldendorf, Germany) and specific primer pairs in a MIC-RUN quantitative PCR system (Bio Molecular Systems, Coomera, Australia). The relative RNA amounts were calculated by the 2^-ΔΔCT^ using 18S RNA as a reference.

### Statistics

Results are presented as mean ± SEM. GraphPad Prism software (v. 9 to 10.4.1) was used to assess statistical significance. Differences between two groups were compared by two-tailed unpaired t-test. Differences between three or more groups were compared by one-way ANOVA followed by the Tukey’s multiple comparisons test. Experiments in which the effects of two variables were tested were analyzed by 2-way ANOVA followed by Tukey’s or Bonferroni’s multiple comparisons test. Differences were considered statistically significant when P < 0.05. Only exact significant P values are reported.

## References

1. Hall IF, Kishta F, Xu Y, Baker AH, Kovacic JC. Endothelial to mesenchymal transition: at the axis of cardiovascular health and disease. Cardiovasc Res. 2024;120:223–236. doi: 10.1093/cvr/cvae021

2. Bischoff J. Endothelial-to-Mesenchymal Transition. Circ. Res. 2019;124:1163–1165. doi: 10.1161/CIRCRESAHA.119.314813

3. Alvandi Z, Bischoff J. Endothelial-mesenchymal transition in cardiovascular disease. Arterioscler Thromb Vasc Biol. 2021;41:2357–2369. doi: 10.1161/ATVBAHA.121.313788

4. Kovacic JC, Dimmeler S, Harvey RP, Finkel T, Aikawa E, Krenning G, Baker AH. Endothelial to mesenchymal transition in cardiovascular disease: JACC state-of-the-art review. J Am Coll Cardiol. 2019;73:190–209. doi: 10.1016/j.jacc.2018.09.089

5. Piera-Velazquez S, Jimenez SA. Endothelial to mesenchymal transition: role in physiology and in the pathogenesis of human diseases. Physiol Rev. 2019;99:1281– 1324. doi: 10.1152/physrev.00021.2018

6. Tombor LS, John D, Glaser SF, Luxán G, Forte E, Furtado M, Rosenthal N, Baumgarten N, Schulz MH, Wittig J, Rogg E-M, Manavski Y, Fischer A, Muhly-Reinholz M, Klee K, Looso M, Selignow C, Acker T, Bibli S-I, Fleming I, Patrick R, Harvey RP, Abplanalp WT, Dimmeler S. Single cell sequencing reveals endothelial plasticity with transient mesenchymal activation after myocardial infarction. Nat Commun. 2021;12:681. doi: 10.1038/s41467-021-20905-1

7. Zeisberg EM, Tarnavski O, Zeisberg M, Dorfman AL, McMullen JR, Gustafsson E, Chandraker A, Yuan X, Pu WT, Roberts AB, Neilson EG, Sayegh MH, Izumo S, Kalluri R. Endothelial-to-mesenchymal transition contributes to cardiac fibrosis. Nat Med. 2007;13:952–961. doi: 10.1038/nm1613

8. Chen P-Y, Qin L, Li G, Wang Z, Dahlman JE, Malagon-Lopez J, Gujja S, Cilfone NA, Kauffman KJ, Sun L, Sun H, Zhang X, Aryal B, Canfran-Duque A, Liu R, Kusters P, Sehgal A, Jiao Y, Anderson DG, Gulcher J, Fernandez-Hernando C, Lutgens E, Schwartz MA, Pober JS, Chittenden TW, Tellides G, Simons M. Endothelial TGF-β signalling drives vascular inflammation and atherosclerosis. Nat Metab. 2019;1:912–926. doi: 10.1038/s42255-019-0102-3

9. Chen P-Y, Qin L, Baeyens N, Li G, Afolabi T, Budatha M, Tellides G, Schwartz MA, Simons M. Endothelial-to-mesenchymal transition drives atherosclerosis progression. J Clin Invest. 2015;125:4514–4528. doi: 10.1172/JCI82719

10. Lu JY, Tu WB, Li R, Weng M, Sanketi BD, Yuan B, Reddy P, Rodriguez Esteban C, Izpisua Belmonte JC. Prevalent mesenchymal drift in aging and disease is reversed by partial reprogramming. Cell. 2025;188:5895–5911.e17. doi: 10.1016/j.cell.2025.07.031

11. Vannahme C, Smyth N, Miosge N, Gösling S, Frie C, Paulsson M, Maurer P, Hartmann U. Characterization of SMOC-1, a novel modular calcium-binding protein in basement membranes. J Biol Chem. 2002;277:37977–37986. doi: 10.1074/jbc.M203830200

12. Choi YA, Lim J, Kim KM, Acharya B, Cho JY, Bae YC, Shin HI, Kim SY, Park EK. Secretome analysis of human BMSCs and identification of SMOC1 as an important ECM protein in osteoblast differentiation. J Proteome Res. 2010;9:2946–2956. doi: 10.1021/pr901110q

13. Awwad K, Hu J, Shi L, Mangels N, Abdel Malik R, Zippel N, Fisslthaler B, Eble JA, Pfeilschifter J, Popp R, Fleming I. Role of secreted modular calcium binding protein 1 (SMOC1) in transforming growth factor β signaling and angiogenesis. Cardiovasc Res. 2015;106:284–294. doi: 10.1093/cvr/cvv098

14. Wang E-L, Zhang J-J, Luo F-M, Fu M-Y, Li D, Peng J, Liu B. Cerebellin-2 promotes endothelial-mesenchymal transition in hypoxic pulmonary hypertension rats by activating NF-κB/HIF-1α/Twist1 pathway. Life Sci. 2023;328:121879. doi: 10.1016/j.lfs.2023.121879

15. Wu Z, Zhu J, Wen Y, Lei P, Xie J, Shi H, Wu R, Lou X, Hu Y. Hmga1-overexpressing lentivirus protects against osteoporosis by activating the Wnt/β-catenin pathway in the osteogenic differentiation of BMSCs. FASEB J. 2023;37:e22987. doi: 10.1096/fj.202300488R

16. Lin Z, Wang C, Wen Z, Cai Z, Guo W, Feng X, Huang Z, Zou R, Fan X, Liu C, Yang H. NEXN regulates vascular smooth muscle cell phenotypic switching and neointimal hyperplasia. JCI Insight. 2025;10. doi: 10.1172/jci.insight.190089

17. Brellier F, Ruggiero S, Zwolanek D, Martina E, Hess D, Brown-Luedi M, Hartmann U, Koch M, Merlo A, Lino M, Chiquet-Ehrismann R. SMOC1 is a tenascin-C interacting protein over-expressed in brain tumors. Matrix Biol. 2011;30:225–233. doi: 10.1016/j.matbio.2011.02.001

18. Gersdorff N, Müller M, Schall A, Miosge N. Secreted modular calcium-binding protein-1 localization during mouse embryogenesis. Histochem Cell Biol. 2006;126:705–712. doi: 10.1007/s00418-006-0200-7

19. Wang Y, Gu J, Du A, Zhang S, Deng M, Zhao R, Lu Y, Ji Y, Shao Y, Sun W, Kong X. SPARC-related modular calcium binding 1 regulates aortic valve calcification by disrupting BMPR-II/p-p38 signalling. Cardiovasc Res. 2022;118:913–928. doi: 10.1093/cvr/cvab107

20. Thomas JT, Eric Dollins D, Andrykovich KR, Chu T, Stultz BG, Hursh DA, Moos M. SMOC can act as both an antagonist and an expander of BMP signaling. Elife. 2017;6. doi: 10.7554/eLife.17935

21. DeGroot MS, Williams B, Chang TY, Maas Gamboa ML, Larus IM, Hong G, Fromme JC, Liu J. SMOC-1 interacts with both BMP and glypican to regulate BMP signaling in C. elegans. PLoS Biol. 2023;21:e3002272. doi: 10.1371/journal.pbio.3002272

22. Gougos A, Letarte M. Primary structure of endoglin, an RGD-containing glycoprotein of human endothelial cells. J Biol Chem. 1990;265:8361–8364

23. Bellón T, Corbí A, Lastres P, Calés C, Cebrián M, Vera S, Cheifetz S, Massague J, Letarte M, Bernabéu C. Identification and expression of two forms of the human transforming growth factor-beta-binding protein endoglin with distinct cytoplasmic regions. Eur J Immunol. 1993;23:2340–2345. doi: 10.1002/eji.1830230943

24. Lastres P, Letamendía A, Zhang H, Rius C, Almendro N, Raab U, López LA, Langa C, Fabra A, Letarte M, Bernabéu C. Endoglin modulates cellular responses to TGF-beta 1. J Cell Biol. 1996;133:1109–1121. doi: 10.1083/jcb.133.5.1109

25. Lastres P, Martín-Perez J, Langa C, Bernabéu C. Phosphorylation of the humantransforming-growth-factor-beta-binding protein endoglin. Biochem. J. 1994;301 ( Pt 3):765–768. doi: 10.1042/bj3010765

26. Xu Y, Kovacic JC. Endothelial to mesenchymal transition in health and disease. Annu Rev Physiol. 2023;85:245–267. doi: 10.1146/annurev-physiol-032222-080806

27. Abouzeid H, Boisset G, Favez T, Youssef M, Marzouk I, Shakankiry N, Bayoumi N, Descombes P, Agosti C, Munier FL, Schorderet DF. Mutations in the SPARC-related modular calcium-binding protein 1 gene, SMOC1, cause Waardenburg anophthalmia syndrome. Am J Hum Genet. 2011;88:92–98. doi: 10.1016/j.ajhg.2010.12.002.

28. Okada I, Hamanoue H, Terada K, Tohma T, Megarbane A, Chouery E, Abou-Ghoch J, Jalkh N, Cogulu O, Ozkinay F, Horie K, Takeda J, Furuichi T, Ikegawa S, Nishiyama K, Miyatake S, Nishimura A, Mizuguchi T, Niikawa N, Hirahara F, Kaname T, Yoshiura K, Tsurusaki Y, Doi H, Miyake N, Furukawa T, Matsumoto N, Saitsu H. SMOC1 is essential for ocular and limb development in humans and mice. Am J Hum Genet. 2011;88:30–41. doi: 10.1016/j.ajhg.2010.11.012

29. Rainger J, van Beusekom E, Ramsay JK, McKie L, Al-Gazali L, Pallotta R, Saponari A, Branney P, Fisher M, Morrison H, Bicknell L, Gautier P, Perry P, Sokhi K, Sexton D, Bardakjian TM, Schneider AS, Elcioglu N, Ozkinay F, Koenig R, Mégarbané A, Semerci CN, Khan A, Zafar S, Hennekam R, Sousa SB, Ramos L, Garavelli L, Furga AS, Wischmeijer A, Jackson IJ, Gillessen-Kaesbach G, Brunner HG, Wieczorek D, van Bokhoven H, FitzPatrick DR. Loss of the BMP antagonist, SMOC-1, causes ophthalmoacromelic (Waardenburg Anophthalmia) syndrome in humans and mice. PLoS Genet. 2011;7:e1002114. doi: 10.1371/journal.pgen.1002114

30. Willison C, Ramachandran V, Chandler NJ, Hillman S, Ashraf T. A novel SMOC1 pathogenic homozygous variant in a fetus with mesomelia of the lower limbs, micrognathia and hypertelorism and an incidental finding of CYP21A2-related congenital adrenal hyperplasia. Prenat Diagn. 2023;43:1674–1677. doi: 10.1002/pd.6485

31. Delgado Lagos F, Elgheznawy A, Kyselova A, Meyer Zu Heringdorf D, Ratiu C, Randriamboavonjy V, Mann AW, Fisslthaler B, Siragusa M, Fleming I. Secreted modular calcium-binding protein 1 binds and activates thrombin to account for platelet hyperreactivity in diabetes. Blood. 2021;137:1641–1651. doi: 10.1182/blood.2020009405

32. Wang H, Dey KK, Chen P-C, Li Y, Niu M, Cho J-H, Wang X, Bai B, Jiao Y, Chepyala SR, Haroutunian V, Zhang B, Beach TG, Peng J. Integrated analysis of ultra-deep proteomes in cortex, cerebrospinal fluid and serum reveals a mitochondrial signature in Alzheimer’s disease. Mol Neurodegener. 2020;15:43. doi: 10.1186/s13024-020-00384-6

33. Dammer EB, Ping L, Duong DM, Modeste ES, Seyfried NT, Lah JJ, Levey AI, Johnson ECB. Multi-platform proteomic analysis of Alzheimer’s disease cerebrospinal fluid and plasma reveals network biomarkers associated with proteostasis and the matrisome. Alzheimers Res Ther. 2022;14. doi: 10.1186/s13195-022-01113-5

34. Wojtas AM, Dammer EB, Guo Q, Ping L, Shantaraman A, Duong DM, Yin L, Fox EJ, Seifar F, Lee EB, Johnson ECB, Lah JJ, Levey AI, Levites Y, Rangaraju S, Golde TE, Seyfried NT. Proteomic changes in the human cerebrovasculature in Alzheimer’s disease and related tauopathies linked to peripheral biomarkers in plasma and cerebrospinal fluid. Alzheimers Dement. 2024. doi: 10.1002/alz.13821

35. Guo Q, Ping L, Dammer EB, Duong DM, Yin L, Xu K, Shantaraman A, Fox EJ, Golde TE, Johnson EC, Roberts BR, Lah JJ, Levey AI, Seyfried NT. Heparin-enriched plasma proteome is significantly altered in Alzheimer’s disease. Mol Neurodegener. 2024;19:67. doi: 10.1186/s13024-024-00757-1

36. Heo G, Xu Y, Wang E, Ali M, Oh HS-H, Moran-Losada P, Anastasi F, González Escalante A, Puerta R, Song S, Timsina J, Liu M, Western D, Gong K, Chen Y, Kohlfeld P, Flynn A, Thomas AG, Lowery J, Morris JC, Holtzman DM, Perlmutter JS, Schindler SE, Vilor-Tejedor N, Suárez-Calvet M, García-González P, Marquié M, Fernández MV, Boada M, Cano A, Ruiz A, Zhang B, Bennett DA, Benzinger T, Wyss-Coray T, Ibanez L, Sung YJ, Cruchaga C. Large-scale plasma proteomic profiling unveils diagnostic biomarkers and pathways for Alzheimer’s disease. Nat Aging. 2025;5:1114–1131. doi: 10.1038/s43587-025-00872-8

37. Oehl-Jaschkowitz B, Vanakker OM, Paepe A de, Menten B, Martin T, Weber G, Christmann A, Krier R, Scheid S, McNerlan SE, McKee S, Tzschach A. Deletions in 14q24.1q24.3 are associated with congenital heart defects, brachydactyly, and mild intellectual disability. Am J Med Genet A. 2014;164A:620–626. doi: 10.1002/ajmg.a.36321

38. Sherva R, Miller MB, Lynch AI, Devereux RB, Rao DC, Oberman A, Hopkins PN, Kitzman DW, Atwood LD, Arnett DK. A whole genome scan for pulse pressure/stroke volume ratio in African Americans: the HyperGEN study. Am J Hypertens. 2007;20:398–402. doi: 10.1016/j.amjhyper.2006.10.001

39. Li J, Yao Y-X, Yao P-S. Circulating biomarkers and risk of hypertension: a two-sample mendelian randomisation study. Heart Lung Circ. 2023;32:1434–1442. doi: 10.1016/j.hlc.2023.09.003

40. Thomas JT, Canelos P, Luyten FP, Moos, M., Jr. Xenopus SMOC-1 Inhibits bone morphogenetic protein signaling downstream of receptor binding and is essential for postgastrulation development in Xenopus. J Biol Chem. 2009;284:18994–19005. doi: 10.1074/jbc.M807759200

41. Wang Y, Wu X. SMOC1 silencing suppresses the angiotensin II-induced myocardial fibrosis of mouse myocardial fibroblasts via affecting the BMP2/Smad pathway. Oncol Lett. 2018;16:2903–2910. doi: 10.3892/ol.2018.8989

42. Dreieicher E, Beck KF, Lazaroski S, Boosen M, Tsalastra-Greul W, Beck M, Fleming I, Schaefer L, Pfeilschifter J. Nitric oxide inhibits glomerular TGF-β signaling via SMOC-1. J Am Soc Nephrol. 2009;20:1963–1974. doi: 10.1681/ASN.2008060653

43. Ji Y, Yan T, Zhu S, Wu R, Zhu M, Zhang Y, Guo C, Yao K. The integrative analysis of competitive endogenous RNA regulatory networks in coronary artery disease. Front Cardiovasc Med. 2021;8:647953. doi: 10.3389/fcvm.2021.647953

44. Ryu J, Choe N, Kwon D-H, Shin S, Lim Y-H, Yoon G, Kim JH, Kim HS, Lee I-K, Ahn Y, Park WJ, Kook H, Kim Y-K. Circular RNA circSmoc1-2 regulates vascular calcification by acting as a miR-874-3p sponge in vascular smooth muscle cells. Mol Ther Nucleic Acids. 2022;27:645–655. doi: 10.1016/j.omtn.2021.12.031

45. Peckert-Maier K, Royzman D, Langguth P, Marosan A, Strack A, Sadeghi Shermeh A, Steinkasserer A, Zinser E, Wild AB. Tilting the balance: therapeutic prospects of CD83 as a checkpoint molecule controlling resolution of inflammation. Int J Mol Sci. 2022;23. doi: 10.3390/ijms23020732

46. Hall IF, Aikawa E, Sluimer J, Baker AH, Kovacic JC. Endothelial to mesenchymal transition in cardiovascular diseases: molecular insights and clinical perspectives. Eur Heart J. 2025. doi: 10.1093/eurheartj/ehaf670

47. Jang Y-S, Choi I-H. Contrasting roles of different endoglin forms in atherosclerosis. Immune Netw. 2014;14:237–240. doi: 10.4110/in.2014.14.5.237

48. López-Novoa JM, Bernabeu C. The physiological role of endoglin in the cardiovascular system. Am J Physiol Heart Circ Physiol. 2010;299:H959–74. doi: 10.1152/ajpheart.01251.2009

49. Zhang X, Zhang Y, Jia Y, Qin T, Zhang C, Li Y, Huang C, Liu Z, Wang J, Li K. Bevacizumab promotes active biological behaviors of human umbilical vein endothelial cells by activating TGFβ1 pathways via off-VEGF signaling. Cancer Biol Med. 2020;17:418–432. doi: 10.20892/j.issn.2095-3941.2019.0215

50. Yan X-C, Cao J, Liang L, Wang L, Gao F, Yang Z-Y, Duan J-L, Chang T-F, Deng S-M, Liu Y, Dou G-R, Zhang J, Zheng Q-J, Zhang P, Han H. miR-342-5p Is a Notch downstream molecule and regulates multiple angiogenic pathways including Notch, vascular endothelial growth factor and transforming growth factor β signaling. J Am Heart Assoc. 2016;5. doi: 10.1161/JAHA.115.003042

51. Adzraku SY, Cao C, Zhou Q, Yuan K, Hao X, Li Y, Yuan S, Huang Y, Xu K, Qiao J, Ju W, Zeng L. Endothelial Robo4 suppresses endothelial-to-mesenchymal transition induced by irradiation and improves hematopoietic reconstitution. Cell Death Dis. 2024;15:159. doi: 10.1038/s41419-024-06546-4

52. Sicklinger F, Zhang Y, Lavine KJ, Simon N, Bucher V, Jugold M, Lehmann L, Konstandin MH, Katus HA, Leuschner F. A minimal-invasive approach for standardized induction of myocardial infarction in mice. Circ. Res. 2020;127:1214–1216. doi: 10.1161/CIRCRESAHA.120.317794

53. Zacchigna S, Paldino A, Falcão-Pires I, Daskalopoulos EP, Dal Ferro M, Vodret S, Lesizza P, Cannatà A, Miranda-Silva D, Lourenço AP, Pinamonti B, Sinagra G, Weinberger F, Eschenhagen T, Carrier L, Kehat I, Tocchetti CG, Russo M, Ghigo A, Cimino J, Hirsch E, Dawson D, Ciccarelli M, Oliveti M, Linke WA, Cuijpers I, Heymans S, Hamdani N, Boer M de, Duncker DJ, Kuster D, van der Velden J, Beauloye C, Bertrand L, Mayr M, Giacca M, Leuschner F, Backs J, Thum T. Towards standardization of echocardiography for the evaluation of left ventricular function in adult rodents: a position paper of the ESC Working Group on Myocardial Function. Cardiovasc Res. 2021;117:43– 59. doi: 10.1093/cvr/cvaa110

54. Fleming I, Fisslthaler B, Dixit M, Busse R. Role of PECAM-1 in the shear-stress-induced activation of Akt and the endothelial nitric oxide synthase (eNOS) in endothelial cells. J Cell Sci. 2005;118:4103–4111. doi: 10.1242/jcs.02541

55. Busse R, Lamontagne D. Endothelium-derived bradykinin is responsible for the increase in calcium produced by angiotensin-converting enzyme inhibitors in human endothelial cells. Naunyn-Schmiedeberg’s Arch Pharmacol. 1991;344:126–129. doi: 10.1007/bf00167392

56. World Medical Association. Declaration of Helsinki: ethical principles for medical research involving human subjects. JAMA. 2013;310:2191–2194. doi: 10.1001/jama.2013.281053

57. Hu J, Popp R, Frömel T, Ehling M, Awwad K, Adams RH, Hammes H-P, Fleming I. Müller glia cells regulate Notch signaling and retinal angiogenesis via the generation of 19,20-dihydroxydocosapentaenoic acid. Journal of Experimental Medicine. 2014;211:281–295. doi: 10.1084/jem.20131494

